# High-salt diet modulates endocrine regulation between cortisol and FGF23

**DOI:** 10.1101/2025.08.14.670453

**Authors:** Matthias B. Moor, Klaudia Kopper, Antoinette Pechère-Bertschi, Michael S. Sagmeister, Rowan S Hardy, Eric Feraille, Daniel G. Fuster, Johannes Loffing, Ganesh Pathare

## Abstract

Excessive dietary salt intake is a global health concern, affecting cardiovascular, renal, and bone health. While the renin-angiotensin-aldosterone system (RAAS) is a known regulator of dietary salt-induced hormonal responses, the impact of adrenal cortisol remains unclear. Here, we performed a retrospective analysis in individuals (n=292) consuming a random diet. Dietary salt intake positively correlated with urinary cortisol and inversely correlated with plasma fibroblast growth factor 23 (FGF23), a bone-derived hormone regulating phosphate and vitamin D homeostasis. Controlled salt diets in healthy individuals confirmed a dose-dependent increase in urinary cortisol and suppression of plasma FGF23. In mice, oral corticosterone, a cortisol analogue, reduced circulating FGF23 levels. RNA-seq analysis of corticosterone-treated MC3T3 osteoblasts identified suppression of FGF23 via glucocorticoid receptor activation, anti-inflammatory pathways, and reduced osteoblast activity. Our findings reveal a novel endocrine cascade where high salt intake elevates cortisol and suppresses FGF23, with potential implications for bone, kidney, and cardiovascular health.

**SIGNIFICANCE STATEMENT:** Excessive dietary salt intake is a global health concern with poorly understood hormonal consequences beyond the renin-angiotensin-aldosterone system. Here, we identify a novel endocrine cascade in which high salt intake elevates cortisol signaling and suppresses fibroblast growth factor 23 (FGF23), a bone-derived hormone central to phosphate and vitamin D homeostasis. These findings are supported by a human cohort on random diets, a controlled dietary salt intervention, and corticosterone experiments in mice and osteoblasts. Mechanistically, cortisol suppresses FGF23 via glucocorticoid receptor activation, anti-inflammatory signaling, and repression of osteoblast activity. These findings have potential implications for bone, kidney, and cardiovascular health, and suggest that dietary salt intake may influence the clinical interpretation of cortisol and FGF23 measurements.

## INTRODUCTION

Excessive dietary salt intake, a hallmark of the Western diet, is a global health concern associated with cardiovascular disease, bone loss, and autoimmune disorders, contributing significantly to morbidity and mortality^1–3^. Beyond its well-established effects on blood pressure, high salt intake disrupts metabolic signaling and hormonal responses^3^. The renin-angiotensin-aldosterone system (RAAS) is the primary hormonal pathway influenced by dietary salt^4^. Notably, adrenal-derived cortisol is known for its role in stress response and immunosuppression, but it may also contribute to salt homeostasis^5,6^. However, the mechanisms underlying this interaction and its impact on the pathophysiology of high salt intake are not yet fully understood.

Fibroblast growth factor 23 (FGF23), a bone-derived hormone, is central to phosphate and vitamin D homeostasis^7^. Elevated FGF23 is implicated in cardiovascular and kidney diseases, while its deficiency causes aging-like phenotypes in animal models^8–10^. Recent studies suggest a link between sodium homeostasis and circulatory FGF23 levels in humans and mice, but the mechanisms remain unclear^11–13.^. Given cortisol’s emerging role in salt homeostasis, and its established effects on bone remodeling through glucocorticoid receptors (GR), it is crucial to investigate the relationship between dietary salt, cortisol, and FGF23.

FGF23 exerts its biological effects by binding to FGF receptors (FGFRs) in the presence of the co-receptor Klotho, a transmembrane protein predominantly expressed in the kidney^14^. Klotho abundance is regulated by physiological stimuli including osmotic stress and dehydration^15^. Distal tubular Klotho governs calcium reabsorption, while proximal tubular Klotho regulates phosphate homeostasis and prevents aging-like phenotypes in mice^14^. Sodium transport regulatory proteins along the nephron are closely linked to FGF23-Klotho axis, and disruption of tubular sodium handling consistently alters circulating FGF23 levels^11,16–19^. In addition, extracellular sodium directly regulates FGF23 formation in osteoblasts^20^. Whether dietary salt and cortisol influence this broader FGF23-Klotho regulatory network remains unknown.

Here, we hypothesize that dietary salt intake modulates endocrine regulation between cortisol and FGF23. To test this, we analyzed human cohort data and data from healthy individuals with and without dietary salt intervention and performed mechanistic experiments in mice and osteoblast cell lines. Our findings reveal a previously unrecognized endocrine cascade, suggesting that dietary salt elevates cortisol signaling, which may contribute to FGF23 suppression.

## MATERIALS AND METHODS

### Human cohort analysis

We retrospectively analyzed data from the BKSR^21^, a single-center observational cohort consisting of individuals. Plasma C-terminal FGF23 (cFGF23) levels were quantified using ELISA. Steroids in urinary samples were quantified using mass spectrometric analysis on a gas chromatograph 7890A coupled to a mass-selective Hewlett-Packard 5975C detector. We analyzed data regarding urinary and serum parameters from 292 individuals. Associations between urinary sodium, cortisol/cortisone, and plasma FGF23 levels were assessed using generalized linear models, after square-root or log transformation as appropriate based on residuals evaluation. Adjustments were made for age, sex, estimated glomerular filtration rate (creatinine CKD-EPI) and body mass index.

### Dietary salt intervention study

Urine and serum samples were employed from previous prospective, crossover-controlled study comprising healthy male volunteers^22^. Participants underwent three 7-day dietary interventions: low-salt diet (LSD, 3 g NaCl/day), normal-salt diet (NSD, 6 g NaCl/day), and high-salt diet (HSD, 15 g NaCl/day), with constant potassium intake. Serum and 24-h urine samples were collected after each dietary phase. Concentrations of urinary free cortisol were measured using an ELISA (Diametra), whereas iFGF23 and cFGF23 levels were measured using an ELISA (Quidel). Changes in plasma hormone levels and urinary parameters across the dietary interventions were compared using paired t-tests (n=12).

### Mouse experiments

C57Bl6 male mice were housed in standard conditions with ad lib access to standard chow and water. All animal experiments were performed in accordance with the guidelines and regulations approved by the University of Birmingham. Drinking water was supplemented with either corticosterone (100 μg/mL, 0.6% ethanol), or vehicle (0.6% ethanol), with an average consumption of 1.25 mg per day. Mice were sacrificed at the end of 2 weeks’ treatment. Plasma iFGF23 and cFGF23 levels were measured using ELISA (Quidel). Control mice received vehicle treatment. Data were analyzed using unpaired t-tests.

### *In vitro* studies

MC3T3-E1 subclone four mouse preosteoblast cells were cultured and differentiated as previously described^20^. Osteogenic differentiation was initiated by culturing the cells in a medium containing 50 μg/ml ascorbic acid and 10 mM β-glycerophosphate for 6 days. Upon differentiation, cells were treated with corticosterone or hydrocortisone (0–200 nM) for 24 hours as noted in results section. *Fgf23* mRNA levels were quantified using qPCR. FGF23 protein levels in the supernatant were measured by ELISA after concentrating the cell supernatant as described earlier^20^. For GR inhibition studies, cells were pre-treated with 50 nM RU-486 (Biotechne) for 1 hour, followed by corticosterone treatment. All experiments were performed in triplicate.

### RNA-seq analysis

Differentiated MC3T3 cells were treated with 100 nM corticosterone for 24 hours (n=3 biological replicates). RNA was extracted, and sequencing was performed commercially at Novogene using the Illumina platform^20^. Differential gene expression (DEG) analysis was conducted using DESeq2, with adjusted p-values (p_adj_ < 0.05) and log_2_ fold-change > 1 considered significant. Gene ontology (GO) analysis was performed to identify enriched biological pathways as previously described^20^.

### Statistical Analysis

Data were presented as mean ± SEM. Statistical significance was determined using t-tests for two-group comparisons. Correlation analyses were conducted using Pearson’s correlation coefficient. All analyses were performed using GraphPad Prism or RStudio.

## RESULTS

### 1. High salt intake in humans activates cortisol signaling and suppresses FGF23 formation

To investigate the association between dietary sodium intake and cortisol signaling, we reanalyzed data from the thoroughly phenotyped population of the Bern Kidney Stone Registry (BKSR), a single-center observational cohort of kidney stone formers^21^. We found that 24-hour urinary sodium, a reliable surrogate marker of dietary sodium intake, was associated with 24-h urinary cortisol (Fig. 1A). Further, urinary sodium was negatively associated with plasma FGF23 levels (Fig. 1B). Notably, urinary cortisone and cortisol tended to show an inverse relationship with plasma FGF23; however, only the association with urinary cortisone but not cortisol reached statistical significance (Fig. 1C, D). After adjusting the models for renal function, age, sex and body mass index in multivariable regression, the association with sodium remained whereas the association with cortisone was attenuated and did not remain significant (p=0.12) (Figures 1E-F).

**Fig. 1:**
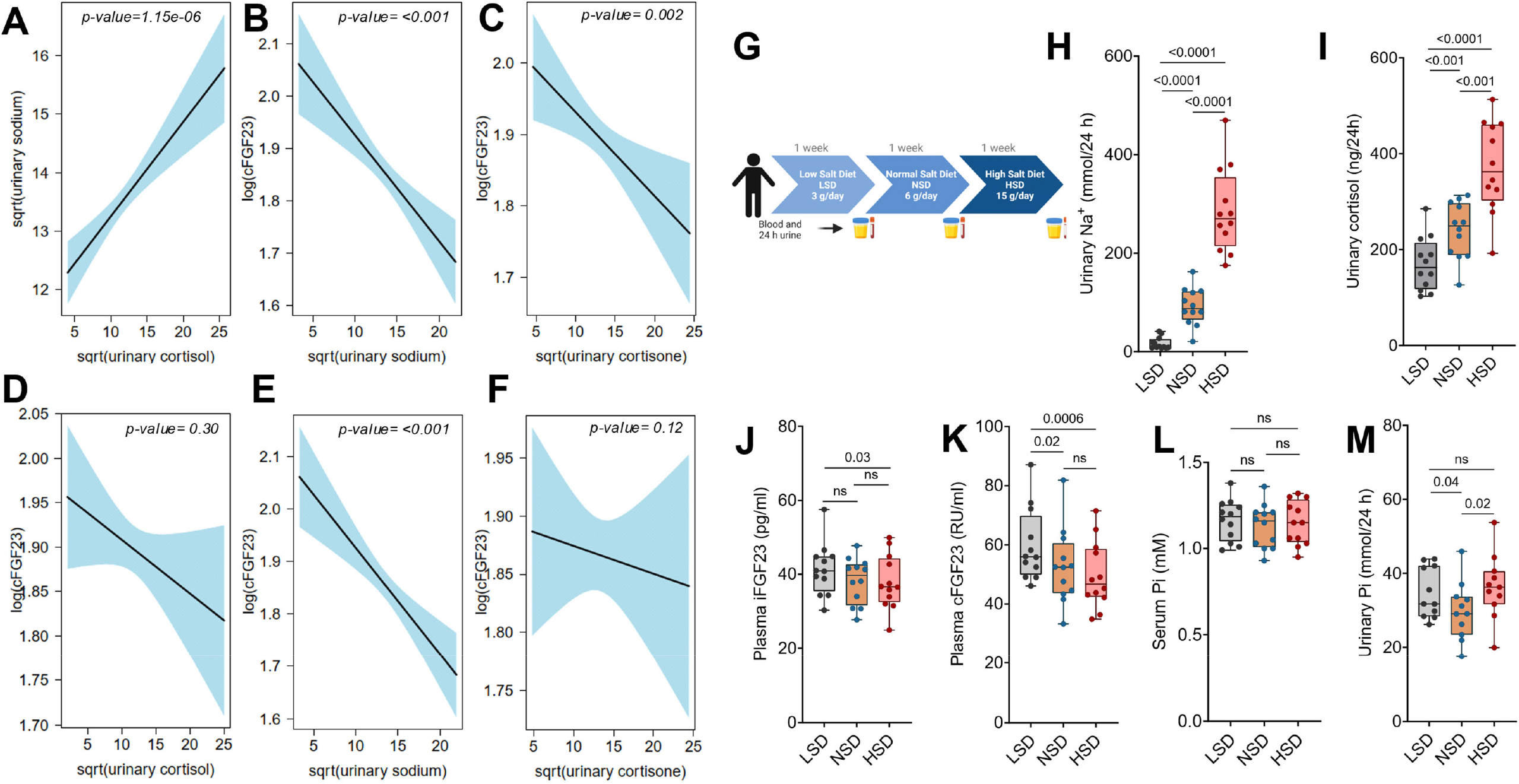
High salt intake in humans elevates cortisol signaling and suppresses FGF23 formation. (A) Positive correlation between 24-hour urinary sodium excretion (a marker of dietary salt intake) and urinary cortisol levels in the BKSR cohort. (B) Inverse correlation between 24-hour urinary sodium excretion and plasma cFGF23 levels in the BKSR cohort. (C) Inverse correlation between plasma cFGF23 levels and 24-hour urinary cortisone levels in the BKSR cohort. (D) Inverse correlation between plasma cFGF23 levels and 24-hour urinary cortisol levels in the BKSR cohort. The p-value indicates the statistical significance of the observed correlation. n= 292. (E) Multivariable regression analysis showing the association between 24-hour urinary sodium excretion and plasma cFGF23 levels after adjustment for age, sex, eGFR, and BMI in the BKSR cohort. (F) Multivariable regression analysis showing the association between 24-hour urinary cortisone and plasma cFGF23 levels after adjustment for age, sex, eGFR, and BMI in the BKSR cohort (p=0.12). n=292 for panels A–F. (G) Schematic representation of the dietary salt intervention study design. (H) Urinary sodium excretion; (I) urinary cortisol levels after dietary salt intervention (low-salt, normal-salt, and high-salt diets). (J) Plasma iFGF23 levels; (K) plasma cFGF23 levels; (L) plasma phosphate levels after dietary salt intervention. (M) Urinary phosphate excretion after dietary salt intervention. Data are presented as mean ± SEM. Statistical significance is denoted with p-value (paired t-test). n=12 per group.

Next, we analyzed serum and urine samples from a prospective crossover-controlled study involving healthy male volunteers^22^. These participants underwent three 7-day dietary interventions: a low-salt diet (LSD, 3 g NaCl/day), a normal-salt diet (NSD, 6 g NaCl/day), and a high-salt diet (HSD, 15 g NaCl/day), with constant potassium intake (Fig. 1G). Dietary sodium intake dose-dependently increased both 24-hour urinary sodium excretion (Fig. 1H) and cortisol levels (Fig. 1I). Interestingly, dietary sodium also dose-dependently suppressed both intact and cFGF23 levels, with a more pronounced effect observed in assays measuring cFGF23 (Fig. 1J/K). No significant differences in serum or urinary phosphate levels were observed across the dietary interventions (Fig. 1L/M). This dissociation between FGF23 suppression and unchanged phosphate levels suggests that the magnitude of FGF23 reduction may fall below the threshold required to perturb phosphate homeostasis, or that compensatory mechanisms maintain phosphate balance despite reduced FGF23.

### 2. Corticosterone suppresses FGF23 formation by GR activation, anti-inflammation, and repression of osteoblast activity

C57Bl6 wild-type mice were administered corticosterone orally for 2 weeks. At the end of the treatment period, blood samples were collected, and intact FGF23 (iFGF23) and C-terminal FGF23 (cFGF23) levels were measured. We found that corticosterone treatment significantly suppressed both iFGF23 and cFGF23 levels in mice (Fig. 2A, B).

**Fig. 2:**
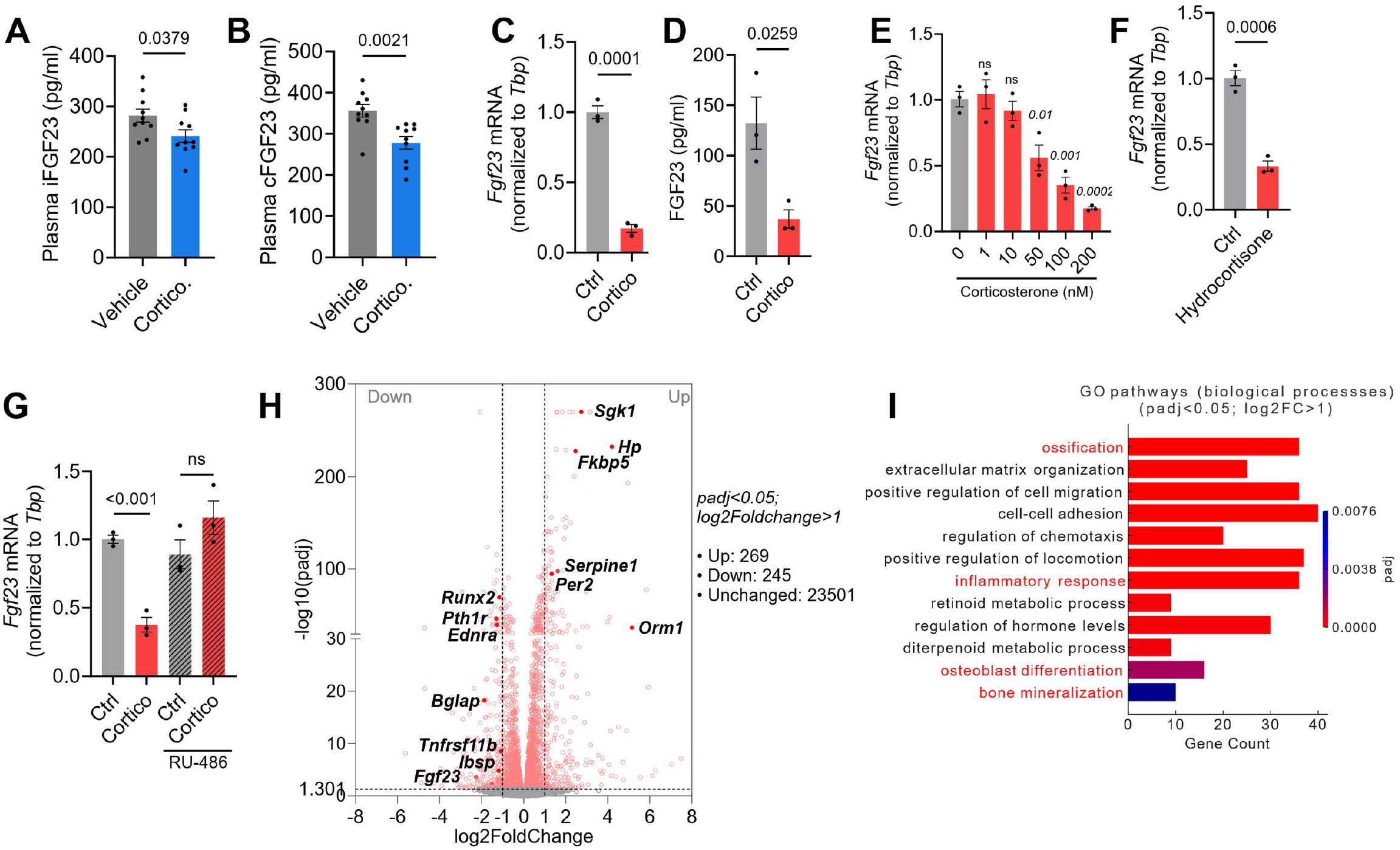
Corticosterone suppresses FGF23 levels via GR activation and repressed osteoblast activity. (A) Plasma iFGF23 levels and (B) cFGF23 levels in C57Bl6 wild-type mice treated with oral corticosterone (n=12 per group). (C) *Fgf23* mRNA expression in cell lysates and (D) FGF23 protein levels in cell supernatant after 100 nM corticosterone treatment for 24 hours in MC3T3 osteoblasts (n=3 per group). (E) Dose-dependent suppression of *Fgf23* mRNA expression in MC3T3 osteoblasts treated with corticosterone for 24 hours (n=3 per group). (F) Suppression of *Fgf23* mRNA expression by 100 nM hydrocortisone after 24-hour treatment (n=3 per group). (G) Rescue of corticosterone (100 nM)-induced suppression of *Fgf23* mRNA expression by pretreatment with RU-486, a glucocorticoid receptor antagonist (n=3 per group). (H) Volcano plot showing differentially expressed genes (DEGs) in corticosterone-treated MC3T3 osteoblasts, highlighting significantly upregulated and downregulated genes (padj < 0.05; |log2FoldChange| > 1). Key genes implicated in GR activation, anti-inflammatory responses, and osteoblast activity are indicated. (I) GO enrichment (biological processes) analysis of DEGs. Pathways related to bone mineralization, inflammatory response, and osteoblast differentiation are highlighted in red. Data are presented as mean ± SEM, with statistical significance denoted with p-value.

To investigate the mechanisms for FGF23 suppression by corticosterone, *in vitro* experiments were conducted using MC3T3 cells. Treatment with 100 nM corticosterone for 24 hours significantly reduced both cFGF23 levels in the cell supernatant and *Fgf23* mRNA expression in cell lysates (Fig. 2C, D). *Fgf23* mRNA expression was dose-dependently suppressed when cells were treated with 0–200 nM corticosterone (Fig. 2E). Treatment with hydrocortisone (100 nM), a synthetic cortisol analogue, also resulted in a reduction in *Fgf23* mRNA expression (Fig. 2F). Pretreatment with RU-486 (GR blocker) completely rescued the corticosterone-induced suppression of *Fgf23* mRNA, suggesting that the effects of corticosterone on *Fgf23* expression are mediated through GR activation (Fig. 2G).

RNA-seq analysis was performed on MC3T3 cells treated with 100 nM corticosterone for 24 hours. Differential gene expression analysis revealed 269 upregulated and 245 downregulated genes (padj < 0.05; log_2_FoldChange > 1), visualized in a volcano plot (Fig. 2H). Gene ontology (GO) analysis of biological processes revealed key pathways, including “ossification,” “inflammatory response,” “osteoblast differentiation,” and “bone mineralization.” (Fig. 2I). Further information is provided in the supplementary datasets 1 and 2 for DEGs and in the complete list of GO pathways. Genes associated with direct corticosteroid signaling, including *Sgk1, Fkbp5*, and *Per1*, were significantly upregulated, highlighting GR activation^*23*^ (Fig. 2H/I). Anti-inflammatory pathways were prominently modulated, with a strong upregulation of classical corticosteroid-responsive genes such as *Hp* (Haptoglobin), *Orm1/2*, and *Serpine1* and a downregulation of *Ednra*^*24*^ (Fig. 2H/I). Simultaneously, genes involved in osteoblast differentiation, such as *Runx2, Pth1r*, and *Ibsp*, were downregulated, indicating reduced osteoblast activity and extracellular matrix remodeling^25^. Suppression of *Bglap* (osteocalcin) indicates reduced osteoblast activity and impaired bone mineralization, while downregulation of *Tnfrsf11b* (OPG), an osteoblast-derived decoy receptor for RANKL, suggests a shift toward enhanced osteoclast-mediated bone resorption (Fig. 2H/I). These findings provide evidence that corticosteroids suppress FGF23 through combined GR signaling, anti-inflammatory effects, and suppression of osteoblast activity (Fig. 2H/I).

## DISCUSSION

Our study identifies a potential endocrine cascade by which high salt intake elevates cortisol signaling and suppresses FGF23 levels, revealing non-classical pathophysiological consequences of high-salt diets on systemic health.

We measured 24-hour urinary cortisol for its accuracy in reflecting systemic cortisol secretion, overcoming the limitations of diurnal variation and inter-individual variability in serum levels^26^. Prior interventional studies with smaller sample sizes suggested that dietary salt intake modulates urinary cortisol levels^5,6^. While real-world dietary sodium variations are less extreme than those in interventional settings, our retrospective analysis showed that even among individuals on random diets, urinary cortisol positively correlates with dietary salt intake. Two possible mechanisms may explain the cortisol rise with high salt intake. First, both aldosterone and cortisol are synthesized in the adrenal cortex, share initial biosynthetic steps, and their levels are affected due to competition for shared precursors and enzymes in the context of salt intake. Suppression of RAAS activity by high salt reduces the demand for aldosterone synthesis, which depends on the enzyme *CYP11B2*. This reduction may divert precursors such as 11-deoxycorticosterone toward cortisol synthesis, potentially increasing cortisol levels^1,27^. Second, a recent study in mice suggest high-salt intake elevates corticosterone levels due to activated hypothalamus-pituitary (HPA) axis and stress response^28^.

We found that high salt-mediated cortisol signaling was negatively associated with plasma FGF23. In the retrospective analysis of individuals consuming random diets, urinary cortisone correlated more strongly with FGF23 than cortisol, likely due to its higher stability. While the association did not remain statistically significant after adjustment, potential residual confounding by renal function cannot be excluded. Notably, in a controlled dietary salt intervention in young adults, high salt intake consistently increased urinary cortisol and suppressed plasma FGF23, supporting this relationship under experimental conditions. The more pronounced suppression of cFGF23 compared to iFGF23 under high-salt conditions may suggest that cortisol modulates not only FGF23 production but also its proteolytic cleavage, though this warrants dedicated investigation.

FGF23 is primarily recognized for its role in phosphate and vitamin D homeostasis, but its connection to salt metabolism is of emerging interest^11–13,16,17^. Aldosterone, a key hormone of the RAAS, directly enhances FGF23 production in bone via mineralocorticoid receptors^11,12^. This was linked to a potential mechanism for FGF23 regulation through dietary salt intervention in previous studies^13^. Our findings suggest that elevated cortisol and reduced aldosterone may collectively contribute to FGF23 suppression in response to high salt intake. However, our study provides limited direct evidence for high salt-induced cortisol suppressing FGF23 levels and highlights the need for further investigation. Furthermore, dedicated *in vivo* experiments are warranted to determine the relative contributions of cortisol and aldosterone to FGF23 regulation under a high-salt diet. Previous studies with potent glucocorticoids dexamethasone and prednisolone have shown both stimulatory and inhibitory effects on FGF23 levels in mice^29,30^. In contrast, our results indicate that the naturally occurring hormone corticosterone suppresses FGF23 production in mice and osteoblast cells via GR activation, anti-inflammatory signaling pathways, and transcriptional repression of osteoblast activity.

The interplay between dietary salt, cortisol signaling, and FGF23 may have important systemic implications. Both high salt intake and elevated cortisol levels contribute to hypertension, autoimmune diseases, inflammation and metabolic syndrome, bone loss and fracture risks, therefore they may create a vicious cycle and amplify the burden of chronic diseases^1,3^. Future studies are warranted to explore pharmacological strategies to counteract glucocorticoid-mediated effects of high-salt diets on bone and cardiovascular health. Furthermore, both cortisol and FGF23 are used in many diagnostic tests^26,31^. Our results suggest that variations in dietary salt intake may influence the interpretations of these measurements.

In conclusion, our study uncovers an endocrine cascade wherein dietary salt intake elevates cortisol signaling, suppressing FGF23 levels. These findings suggest an unconventional role of dietary salt in systemic health and highlight the need for further study.

## Acknowledgments

The authors sincerely thank Monique Carrel for her excellent technical assistance. We are grateful to Prof. Franz Weber (Centre of Dental Medicine, University of Zurich) for providing the MC3T3-E1 cells, and to Prof. Mariusz P. Kowalewski (Institute of Veterinary Anatomy, University of Zurich) for critically reading the manuscript.

## Author contributions

GP conceptualized research; MBM, APB, MSS, RSH, EF, DF and GP designed research; MBM, KK, APB, MS and GP performed research/reanalyzed human cohort data; JL contributed new reagents/analytic tools; MBM, KK, APB, MSS, RSH, EF, DF, JL and GP analyzed the data; GP wrote the manuscript; MBM, KK, APB, MSS, RSH, EF, DF and JL reviewed and edited the manuscript; GP and JL were responsible and funding acquisition; GP was responsible for supervision.

## Data availability

The raw RNA-seq data used in this publication have been deposited in NCBI’s Gene Expression Omnibus and are accessible through accession number GSE304562. All data are available from the authors upon request.

